# Substantial gene flow caused by long-term translocation between wild populations of the Peruvian scallop (*Argopecten purpuratus*) is supported by RAD-Seq analyses

**DOI:** 10.1101/2020.09.09.289470

**Authors:** Ximena Velez-Zuazo, Sergio P. Barahona, Omar G. Melo, Eric Hanschke, Ian Hanschke, Monica C. Santa-Maria

**Author notes:** Correspondence Sergio P. Barahona, Universidad Científica del Sur, Carr. Panamericana Sur 19, Villa El Salvador, Lima, Perú.

## Abstract

The Peruvian scallop (*Argopecten purpuratus*, Lamarck 1989) is a marine bivalve of high commercial value in the aquaculture industry, with wild populations distributing from northern Peru to Chile. Its growing demand in the world aquaculture markets and limited availability of hatchery-based seeds, caused long-term seed translocations among wild populations to recover depleted local populations and for production needs. We investigated long-term translocations effects on the genetic diversity and structure of wild populations using next-generation RAD sequencing. We sampled individuals from Sechura, Lobos de Tierra, Samanco and Bahia Independencia in Peru, and La Rinconada in Northern Chile. We identified 6275 polymorphic RAD tags and 8345 SNPs for the five populations. We estimated high observed heterozygosity for all populations and high SNP frequency compared to similar studies on marine bivalves. We detected no spatial divergence among populations in Peru (pairwise FST ranged from 0 to 0.003), but strong differentiation with the population in Chile. Migration rate estimates suggested asymmetric directionality of seed translocation. Overall, our results support a remnant effect of an intense historic translocation and on-going gene flow among wild populations in Peru, challenging the identification of outlier loci and certification of sustainable origin of cultured scallops using genetic markers.

## 1. INTRODUCTION

The Peruvian scallop (*Argopecten purpuratus*) is the most exported marine bivalve from the Peruvian aquaculture industry and an important species in the global bivalve market (Cavero Cerrato & Rodríguez Pinto, 2008). It has a long planktonic larval stage and inhabits sheltered bays along the coasts of Peru and Chile, from Paita (5.1°S) to Valparaiso (33.1°S) (Wolff & Mendo, 2000; von Brand, Abarca, Merino, & Stotz, 2016). This species has been subject to local translocations for decades, fueled by the expansion in Peruvian scallop harvest during the 1980s (Mendo, Wolff, Carbajal, Gonzáles, & Badjeck, 2008). In 1982-1983 El Niño Southern Oscillation (ENSO) event caused the overgrowth of *A. purpuratus* populations, resulting in overexploitation and jeopardizing the integrity of wild stocks. In response, the Peruvian government promoted the adoption of aquaculture practices to guarantee the sustainability of this resource. But, the limited availability of hatchery-produced seeds led to periodical extractions and translocations of young individuals from healthy wild populations to recover depleted local populations (Mendo et al., 2008). This practice was aimed to promote the recovery of diminished natural stocks and to facilitate scallop production in nearby hatcheries.

Translocation of individuals among wild populations of marine species with commercial value is a widespread practice with consequences that are not yet fully understood (Lemer & Planes, 2012). Translocation is defined as the man-made movement of individuals from a given population to another within the species range of distribution (Beaumont, 2000). This practice serves specific purposes, the main being the recovery of depleted populations for securing harvest (Carlton & Mann, 1996). Evidence indicates that estimates and distribution of genetic diversity and relatedness can change in wild populations as a result of translocations. For example, a genetic homogenization in wild populations of previously distinct black-lip pearl oyster (*Pinctada margaritifera*) was found following ten years of translocations (Arnaud-Haond et al., 2004). A recent study in the same species reported high levels of genetic diversity in wild populations adjacent to farmed populations due to “gene leaking” (Lemer & Planes, 2012).

The intensity and directionality of translocations of Peruvian scallops are poorly documented. An early study investigated the level of genetic structure for three Peruvian scallop populations in Peru (Sechura, Samanco, and Bahia Independencia). They found a significant, albeit weak, population differentiation using nine microsatellite and mitochondrial markers (Marín, Fujimoto, & Arai, 2013). The authors suggested that, despite intense translocations, at least one population retained part of its genetic signature (Bahia Independencia) due to asymmetric translocations and to the effect of geographic distance for realized larval dispersion. A recent study investigated the genetic structure in Peruvian scallop populations spanning its entire geographic distribution using two mitochondrial markers (COI and cytb genes) and found a slightly phylogeographical structure between Peru and Chile populations (Acosta-Jofré et al., 2020). Their results suggested that scallop populations experienced a recent population range expansion, with moderate to low levels of gene flow after the expansion. Both studies attributed the observed patterns to high man-mediated gene flow through seed translocation, and suggested further studies with higher marker resolution to understand population structuring.

Genome-wide molecular markers could provide the resolution power needed to analyze and detect structure in large populations with high gene flow (Waples, 1998). Genome-wide markers, particularly those generated by genotyping-by-sequencing approaches (Davey et al., 2011) such as Restriction-Site Associated DNA (RAD) markers (Baird et al., 2008) can detect variation and resolve fine-scale population structure for species with large dispersal capacity and potential high gene flow, e.g. mosquitoes (Rašić, Filipović, Weeks, & Hoffmann, 2014) and American lobster (Benestan et al., 2015). We evaluated the genome-wide variation in individuals of the Peruvian scallop from five wild populations along their distribution range using thousands of SNPs makers obtained from next-generation RAD sequencing. The objective was to investigate the influence of long-term translocations on the current genetic structure of Peruvian scallop wild populations. We (1) estimated levels of genomic diversity among and within wild populations, (2) described the spatial population structure, and (3) assessed the level and directionality of translocations among wild populations.

Despite its major role in the Peruvian aquaculture industry, the sustainable harvest of Peruvian scallop is not totally guaranteed. To this day, the Peruvian aquaculture industry continues to rely on wild seed extraction, with much of the extracted seed being moved to nearby hatcheries or distant ones adjacent to wild population (E. Hanschke, personal communication). This hinders the performance of Peruvian producers in international markets (I. Hanschke, personal communication). Traceability tools, such as genome-wide molecular markers, could provide a sustainable origin certification of aquaculture species, demanded by the global bivalve market (Nielsen et al., 2012). In this study we also discuss perspectives on the use of genetic markers and population-wide genetic descriptors (i.e. heterozygosity and inbreeding coefficients) for traceability.

## 2. MATERIALS AND METHODS

### 2.1. Sample collection and DNA purification

Between August 2014 and February 2015, we collected 57 individuals of *A. purpuratus* from wild populations covering a distribution ranging from northern Peru (Sechura-SEC, Lobos de tierra-LDT, Samanco-SAM, and Bahia Independencia -IND) to northern Chile (La Rinconada-RIN) (Figure 1, Table 1). We obtained approximately 200 mg of adductor muscle tissue and stored it in RNAlater (Ambion, Thermo Fisher Scientific, Waltham, MA USA) at room temperature for the first 24 hours and then stored permanently at −20°C. Whole genomic DNA was isolated using the GeneJET Genomic DNA Purification Kit (Thermo Fisher Scientific, Waltham, MA USA) following manufacturer instructions.

**FIGURE 1.**
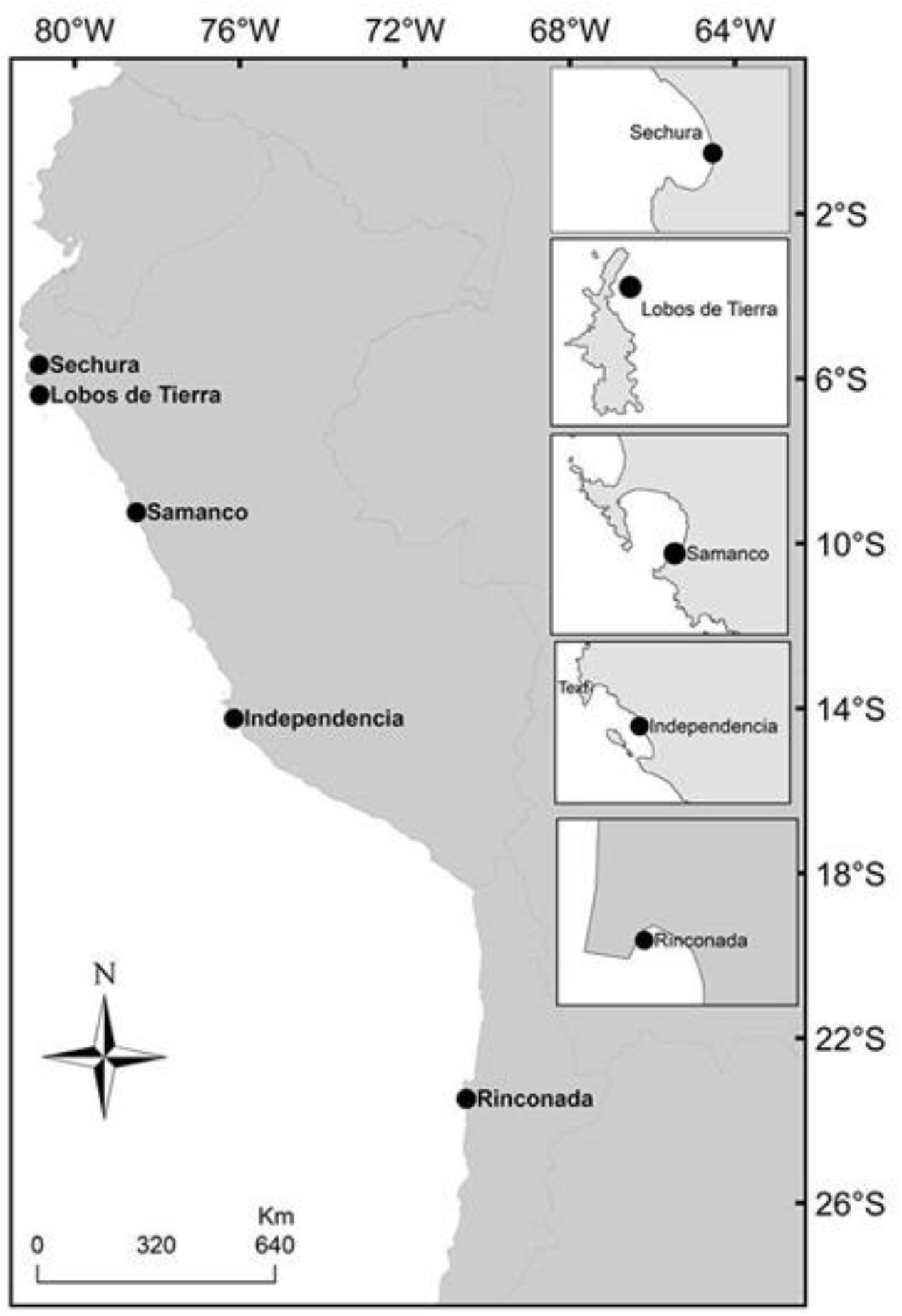
Peruvian scallop (*Argopecten purpuratus*) populations investigated in this study.

**TABLE 1.** Molecular diversity estimations for five populations of bay scallop. N = number of individuals, Ho = observed heterozygosity, He = expected heterozygosity, π = nucleotide diversity, Fis = inbreeding coefficient. Standard deviation is indicated within parenthesis.

| Population | Latitude - Longitude | N | $H_o$ | $H_e$ | $\pi$ | $F_{IS}$ |
| --- | --- | --- | --- | --- | --- | --- |
| Sechura | 5°47'S - 80°55'W | 12 | 0.1855<br>(0.0273) | 0.1814<br>(0.0183) | 0.1897<br>(0.0201) | 0.0285<br>(0.0619) |
| Lobos de Tierra | 6°25'S - 80°50'W | 12 | 0.1924<br>(0.0322) | 0.1862<br>(0.0197) | 0.1950<br>(0.0217) | 0.0311<br>(0.0707) |
| Samanco | 9°11'S - 78°33'W | 12 | 0.1677<br>(0.0224) | 0.1756<br>(0.0174) | 0.1834<br>(0.0190) | 0.0655<br>(0.0767) |
| B. Independencia | 14°14'S - 75°27'W | 9 | 0.2044<br>(0.0264) | 0.2041<br>(0.0171) | 0.2164<br>(0.0193) | 0.0455<br>(0.0752) |
| La Rinconada | 23°27'S - 70°30'W | 12 | 0.1932<br>(0.0271) | 0.1970<br>(0.0193) | 0.2059<br>(0.0211) | 0.0556<br>(0.0790) |

### 2.2. Genomic libraries, denovo assembly and dataset filtering

To construct the Genomic RAD libraries, we followed the original method (Etter, Bassham, Hohenlohe, Johnson, & Cresko, 2011) with minor modifications. Briefly, we digested about 1μg of high-quality and high-concentration genomic DNA with a low frequency restriction enzyme (‘Sbf1’, Thermo Fisher Scientific). We ligated the digested fragments to a modified Solexa adaptor P1 with a unique 5bp barcode and pooled the DNA fragments from individuals bearing unique barcode identifiers for further fragmentation using a COVARIS S220 Focused-ultrasonicator (Woburn, MA, USA). Fragments ranging from 300bp to 600bp were selected from a low melting point agarose gel, ligated to a P2 adapter, and enriched using PCR. RAD-seq libraries were run with a single end 100bp kit in an Illumina HiSeq2500 sequencer using fast mode. Six Illumina HiSeq2500 lanes were sequenced at the University of Chicago Genomics Facility (USA) for this specific study.

We used FASTQ (Andrews, 2010) to assess the quality of RAD sequences. Sequences with Phred quality scores greater than 20 were kept. For sample demultiplexing, denovo assembly, and RAD loci and SNPs identification we used the perl modules implemented in STACKS v1.10 (Catchen et al., 2011; Catchen et al., 2013). For the *denovo* assembly and to generate a catalog of usable loci, we established the following parameters: (1) a minimum depth of six RAD reads per stack or locus (−m), (2) a maximum of two mismatches to add reads (secondary) to a given stack, and (3) a maximum number of three mismatches allowed (−M) between loci within each individual. The resulting genotypes were combined using the *populations* module in Stacks, filtered to include all loci shared for at least 75% of individuals (r = 075 from each population (p = 5) and minimum allele frequency equal or greater than 0.01). To generate final output files for downstream analyses, we used a combination of different output formats such VCF, Genepop, and STRUCTURE (Pritchard, Stephens, & Donnelly, 2000).

### 2.3. Molecular estimates and genetic analyses

We calculated basic molecular estimates at the population level (i.e. observed heterozygosity-Ho, expected heterozygosity-He, and inbreeding coefficient-Fis) using the output provided by the “populations” module from STACKS.

We analyzed deviations from the Hardy-Weinberg equilibrium using the GenoDive software v2.0 (Meirmans & Van Tienderen, 2004). To investigate how genetic diversity is distributed among populations, we conducted a molecular variance analysis (AMOVA) and tested for population structure estimating the Weir and Cockerham’s *F*_ST_ unbiased estimator (Weir & Cockerham, 1984) between all population pairs. We used GenoDive with 999 permutations and a Bonferroni correction (Rice, 1989) to adjust p-values for these calculations. To further test for significant structure among populations, we conducted a Principal Components Analysis (PCA) using the package Adegenet version 2.0.1 (Jombart, 2008; Jombart & Ahmed, 2011). This package was implemented in the software R within Rstudio 1.0.136 (Rstudio Team, 2015).

To investigate the level and direction of historical translocations between wild populations, we estimated the number of migrants (Nm) among the wild populations using a coalescent approach implemented in the program Migrate-n v3.6.4 (Beerli & Felsenstein, 2001). Migrate-n uses Bayesian inference to estimate effective population size (θ parameter) and migration rates (M parameter), which can be used to calculate Nm (Nm = Mxθ). Migrate-n was run with the full migration Matrix model, adjusting the transition/ transversion rate value to 2.0, for nuclear DNA. The dataset for this analysis included loci that were shared by all individuals from all populations, excluding all candidate markers suggested to be under positive selection by Bayescan (Foll & Gaggiotti, 2008).

## 3. RESULTS

We developed and sequenced six genomic RAD libraries that yielded approximately 260 million sequences. On average, <1% of these sequences had ambiguous barcodes, <1% were low quality sequences, and less than 7% had ambiguous RAD-tags. These estimates varied among individuals (see Supplementary Data 1). After the initial screening and filtering, 225,410,859 sequences were retained for further analyses. The number of retained sequences varied among individuals and populations. The number of sequences per individual varied between 498,875 and 9,743,838 (mean value: 3,954,576.47, SD± 2,101,529.36) and the mean percentage of retained sequences was 85% (see Supplementary Data 2). The average Phred score value was 30. All sequences were pruned to 80bp to minimize variability in the sequence quality. The denovo assembly identified 9,593 RAD loci. After filtering the data to obtain only polymorphic loci for at least 70% of individuals from each population, we recovered 8,354 polymorphic RAD loci (with ≥1 SNP) and 24,218 SNPs. The number of SNPs was further reduced to 7,608 when including only loci shared by all individuals from all populations (zero missing data) and removing outliers SNPs (SNP-outliers = 4) for migration estimates.

### 3.1. Genetic diversity

Observed heterozygosity (Ho) varied between 0.17 and 0.20 (Table 1). The population of Independencia Bay exhibited the highest observed heterozygosity (Ho = 0.20) while Samanco exhibited the lowest value (Ho = 0.17). Similarly, individuals from Bahia Independencia presented the highest nucleotide diversity (π = 0.22) while Samanco exhibited the lowest diversity (π = 0.1931). The population from Samanco exhibited the highest estimate for the inbreeding coefficient (Fis = 0.085) while Lobos de Tierra and Sechura had the lowest (Fis = 0.03) (Table 1).

### 3.2. Population structure and gene flow analysis

Molecular variance analysis indicated that most of the genetic diversity is distributed within individuals (92.6%, *F*_IT_ = 0.74) and to a lesser extent among individuals (6.3%, *F*_IS_ = 0.064, *P* = 0.001) and among populations (1.1%, *F*_ST_ = 0.011, *P* = 0.001). Differentiation values among populations (*F*_ST_) varied between 0 and 0.032 (Table 2). Significant genetic differentiation was observed between the population of La Rinconada (Chile) and the other populations (average *F*_ST_ = 0.029, *P* = 0.001). The lowest values of genetic differentiation were observed between Samanco compared to Bahia Independencia and Lobos de Tierra. The spatial structure detected between La Rinconada and the other populations was further observed in the principal component analysis (Figure 2).

**TABLE 2.**
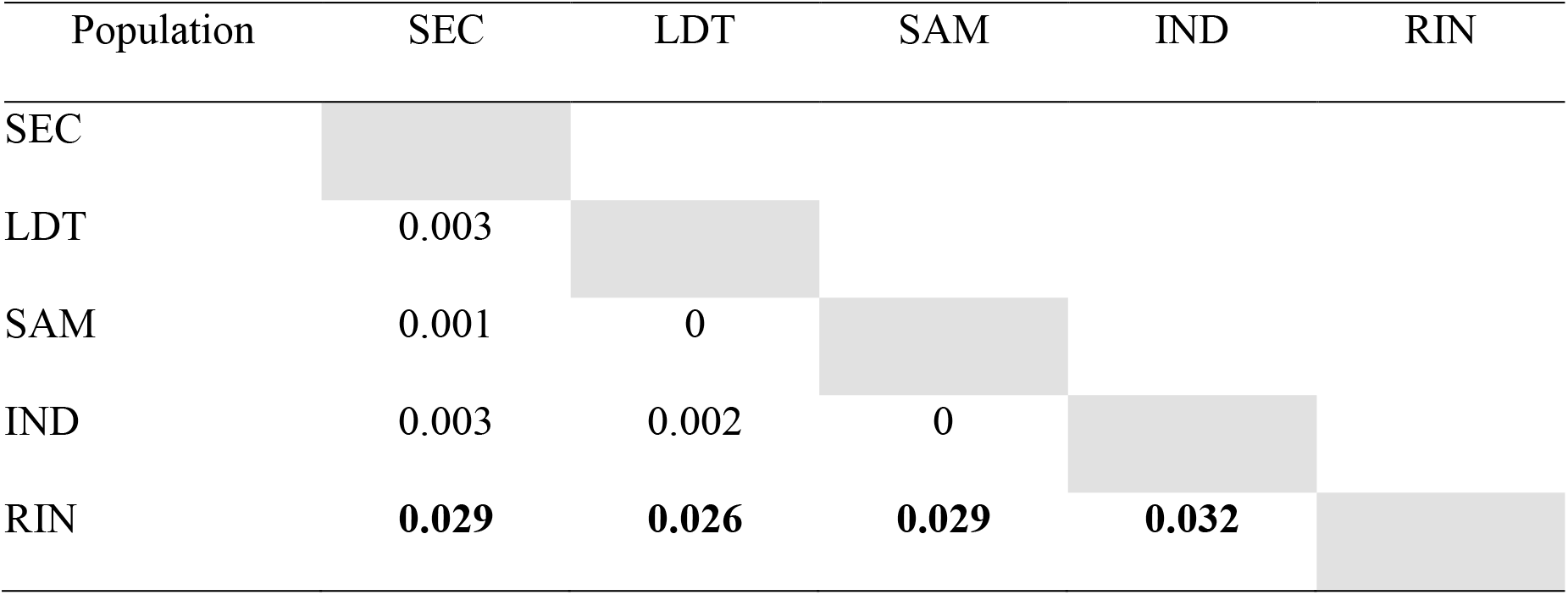
Triangular matrix with population differentiation estimations among five bay scallop’s natural beds populations. F_ST_ values can be observed under the diagonal. Significant values are in bold. SEC = Sechura, LDT = Lobos de Tierra, SAM = Samanco, IND = Bahia Independencia, and RIN = La Rinconada.

| Population | SEC | LDT | SAM | IND | RIN |
| --- | --- | --- | --- | --- | --- |
| SEC |  |  |  |  |  |
| LDT | 0.003 |  |  |  |  |
| SAM | 0.001 | 0 |  |  |  |
| IND | 0.003 | 0.002 | 0 |  |  |
| RIN | **0.029** | **0.026** | **0.029** | **0.032** |  |

**FIGURE 2.**
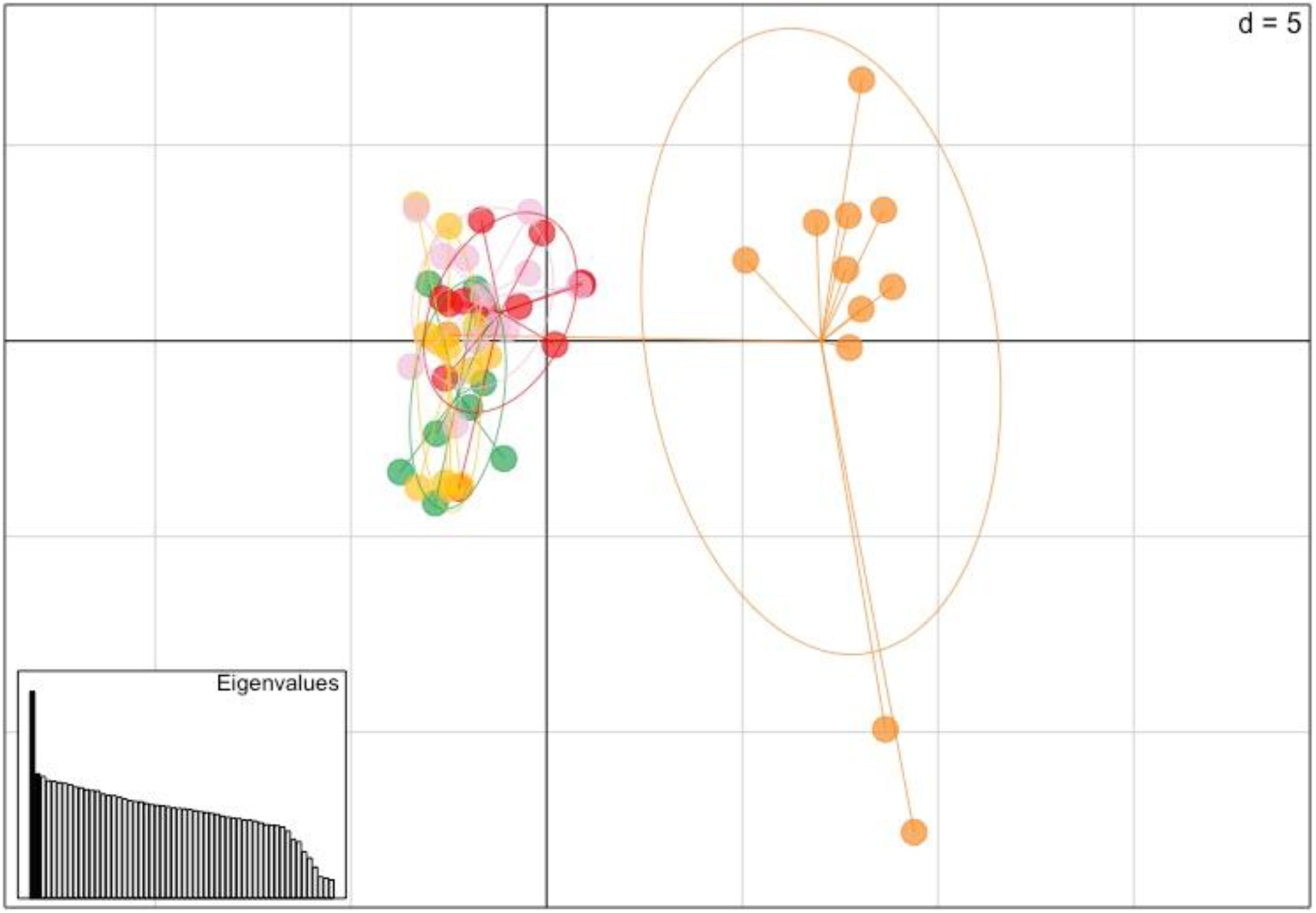
Differentiation at neutral loci among five wild populations of *A. purpuratus* as suggested by principal components analysis. Populations are color-coded: Sechura (pink), Lobos de Tierra (green), Samanco (yellow), Bahia Independencia (red), and La Rinconada (Chile) (orange). The three axes from the principal component analyses accounted for 9.2% of the genetic variation.

Gene flow analysis implemented in Migrate-n was carried out using 3027 loci shared among the five wild populations. In average, considering all possible directions, northward gene flow was higher compared to southward gene flow (Nm_Northward_ = 6848.17, Nm_Southward_ = 2524.63). The analysis showed asymmetrical gene flow between Chilean and Peruvian populations: a strong gene flow from La Rinconada to the Peruvian populations (mean Nm = 14855.4) and a weak gene flow from the Peruvian populations to La Rinconada (mean Nm = 1946.45). Within the Peruvian populations, high gene flow from Lobos de Tierra to the other populations was observed (mean Nm = 6086.25), while low gene flow was detected among the other populations (Sechura, Samanco and Bahia Independencia) (Table 3).

**TABLE 3.** Mean estimates of mutation-scaled effective population sizes per population (θ), migration rate (M) and number of migrants per generation (Nm) for each and migration directionality, calculated by Migrate-n. Nm was calculated as Mxθ.

| **Population** | **Code** | **$\theta$** |  |
| --- | --- | --- | --- |
| Sechura | SEC | 1.5 |  |
| Lobos de Tierra | LDT | 4.5 |  |
| Samanco | SAM | 0.5 |  |
| B. Independencia | IND | 0.5 |  |
| La Rinconada | RIN | 11.5 |  |
| **Migration directionality** | **M** | **Nm** | **Movement** |
| SEC → LDT | 835.8 | 1253.70 | Southward |
| SEC → SAM | 1137.5 | 1706.25 | Southward |
| SEC → IND | 1073.6 | 1610.40 | Southward |
| SEC → RIN | 681.8 | 1022.70 | Southward |
| LDT → SEC | 1393.3 | 6269.85 | Northward |
| LDT → SAM | 1378.7 | 6204.15 | Southward |
| LDT → IND | 1337.4 | 6018.30 | Southward |
| LDT → RIN | 1300.6 | 5852.70 | Southward |
| SAM → LDT | 1148.1 | 574.05 | Northward |
| SAM → SEC | 1079.5 | 539.75 | Northward |
| SAM → IND | 1335.4 | 667.70 | Southward |
| SAM → RIN | 952.9 | 476.45 | Southward |
| IND → LDT | 1100.2 | 550.10 | Northward |
| IND → SEC | 1217.1 | 608.55 | Northward |
| IND → SAM | 1035.5 | 517.75 | Northward |
| IND → RIN | 867.9 | 433.95 | Southward |
| RIN → LDT | 1320.9 | 15190.35 | Northward |
| RIN → SEC | 1208.1 | 13893.15 | Northward |
| RIN → SAM | 1344.1 | 15457.15 | Northward |
| RIN → IND | 1294 | 14881.00 | Northward |

## 4. DISCUSSION

Despite being the most economically valuable bivalve aquaculture species in Peru, up to date it is impossible to differentiate batches of exported wild Peruvian scallops from those produced in a sustainable way. This is mostly due to the lack of a reliable regulation system and/or methods for traceability and certification of origin for hatchery produced batches. Genetic tools offer an opportunity for reliable traceability of seafood (Bernatchez et al., 2017) given detailed information on the genetic structure and distribution of the source populations is available. Two previous studies used mitochondrial markers to assess the genetic diversity and population structure in the Peruvian scallop (Marín et al., 2013; Acosta-Jofré et al., 2020). These markers were useful in determining genetic differences between populations, but they lack the power to detect small, but significant, genetic differences in nearby and large populations (Waples, 1998). In our study, we obtained estimates of genetic diversity and assessed the degree of population structure exploring the variation in 7608 SNPs.

We found high levels of genetic diversity and SNP frequency compared to other bivalve species. The mean estimates of genetic diversity, as indicated by Ho, was 0.1886 for all wild populations assessed in our study and higher when compared to other marine bivalves. For example, a mean Ho of 0.1085 for wild populations was reported for the black-lip pearl oyster *Pinctada margaritifera* (Lal, Southgate, Jerry, & Zenger, 2016). The observed heterozygosity for *A. purpuratus* was nearly equal to the expected heterozygosity (He, Table 1). This contrasts with other studies reporting heterozygote deficiency as a result of diverse proposed mechanisms (Miller, Versace, Matthews, Montgomery, & Bowie, 2013; Peñaloza, Bishop, Toro, & Houston, 2014; Lal et al., 2016). Similar to the genetic diversity levels, our estimates of SNP frequency for *A. purpuratus* were also higher compared to other bivalve species. We detected, in average, a SNP every 20bp, compared to estimates of 1SNP/30bp for the Chilean mussel and 1 SNP/40bp for the Pacific oyster (Sauvage, Bierne, Lapegue, & Boudry, 2007). Our estimates agree with high polymorphic levels previously reported for bivalves.

Overall, the genetic diversity did not differ significantly among populations, albeit the population of Bahia Independencia which exhibited the highest genetic diversity values (Ho = 0.2044). This agrees with a previous study where the population of Bahia Independencia exhibited higher genetic diversity than populations from Sechura and Samanco (Marín et al., 2013). Sechura and Samanco populations had the lowest genetic diversity estimates, which is in agreement with a previous study (Marín et al 2013). These two populations are adjacent to hatcheries, so the low levels of diversity could be the result of migrants coming from them. A similar scenario has been observed previously for the black-lip pearl oyster (Lal et al., 2016). Low genetic diversity in hatchery populations is expected due to the reproduction dynamics in captivity, where a few progenitors are responsible of all the individuals produced, thus promoting allele loss. This has been observed in populations of captive invertebrates (Miller et al. 2013).

Samanco (Fis = 0.0655) and La Rinconada (Fis = 0.0556) (Table 1) exhibited the highest inbreeding coefficient while Lobos de Tierra and Sechura exhibited the lowest values. The surplus of homozygotes can be explained by self-fertilization caused by closed circulation in these bays. The Peruvian scallop is a functional hermaphrodite with external cross-fertilization and sperm release before the oocytes, to prevent self-fertilization (Uriarte, Rupp, & Abarca, 2001). And the areas in Samanco and La Rinconada present gyres or gyre-like currents (Avendaño D., Cantillánez S., Thouzeau, & Peña, 2007; Tenorio, 2016) which promote larval retention. In contrast, the lower inbreeding values detected in Lobos de Tierra Islands and Sechura Bay (Table 1) may be due to low larval retention in these areas (Flores-Valiente et al., 2019).

Population structure across the distribution range of *A. purpuratus* was detected between La Rinconada (Chile) and the Peruvian populations. Using the loci present in >= 70% of the individuals of all populations we were able to distinguish La Rinconada population from the rest. This agrees with Acosta-Jofré et al (2020), who also detected significant genetic differentiation between Antofagasta’s population (very close to La Rinconada) with the Peruvian populations. La Rinconada population is located far from the other populations; the linear distance to the nearest wild population in Peru is nearly 1250 km, making it very unlikely to be naturally colonized by larval dispersion from other areas (Acosta-Jofré, 2020). The differentiation was further supported with the ordination analysis, where La Rinconada population clearly distinguishes from the rest (Figure 2). Within Peru, we did not detect significant differentiation among wild populations (Table 4). Pairwise *F*_ST_ values ranged from 0 (IND vs SAM, LDT vs SAM) to 0.003 (LDT vs SEC, IND vs SEC). Similarly, the molecular variance analysis indicates most of the variation (92.6%) exists within, rather than among populations, thus providing further evidence for the weak or lack of population differentiation among all Peruvian wild populations. Considering the geographic distance separating the wild populations in Peru, the high level of genetic admixture is remarkable. Exception are the populations of Sechura and Lobos de Tierra which are separated by less than 150 km. If we account for the moderate dispersal capability of the Peruvian scallop and the “weak source (SEC) – strong sink (LDT)” scenario suggested for the populations of Sechura and Lobos de Tierra (Flores-Valiente et al., 2019), high levels of genetic admixture are expected. But, for the other populations, the distance between them is >600 km, right in the upper limit of the dispersal distance estimated for species with high larval dispersion capabilities (Van Wyngaarden et al., 2017). Except for Flores-Valiente et al (2019), no other study has assessed the potential role that marine currents in Peru play in larval transport and retention (see Barahona, Velez-Zuazo, Santa-Maria, & Pacheco, 2019). In any case, we would expect to detect population structure at least between Bahia Independencia and the rest of sites. Previous genetic distinction for this population was observed by Marín et al. (2013), who suggested that the geographic and oceanographic characteristics of this bay strongly limited natural gene flow with other populations.

We do not overrule the homogenizing role of larval dispersal facilitated by currents and oceanographic events (Miller et al. 2013), in combination with the large population size and high fecundity of marine bivalves (Cano, Shikano, Kuparinen, & Merilä, 2008). The Peruvian scallop has a moderate pelagic larval duration of 16-25 days in the water column, depending on the water temperature, before settlement (von Brand, Merino, Abarca, & Stotz, 2006). The combination of the northward passive larval displacement caused by the Peruvian Coastal Current (PCC) and the southward displacement caused by the movement of warm water masses during ENSO events could facilitate the bidirectional population connectivity (Barahona et al., 2019). The recolonization of extinct natural beds (Wolff & Mendo, 2000) and the formation of temporal intermediate aggregations (Ysla, 2009) during and after ENSO could increase the genetic connectivity. But, given the geographic peculiarities and large distances separating Peruvian scallop natural beds, larval dispersal capabilities and marine currents alone are unlikely to explain the level of admixing we observed. Our results thus suggest a significant effect of historical translocations in gene-flow and homogenization of wild populations along the Peruvian coast beyond what would be expected by larval dispersion alone.

In any case, we detected an asymmetrical gene flow among the Peruvian and La Rinconada populations. The strong gene flow observed from the Chilean population agrees with the cold northward Humboldt Current (PCC). We found that mean northward gene flow is stronger than mean southward gene flow, which suggests that the effect of ENSO warm periods is weaker compared with the constant flow of the Peruvian Coastal Current for this species. Among Peruvian populations, we found evidence indicating that Lobos de Tierra is a source of migrants, which is consistent with reports of that location being a common source of seeds for translocations (Mendo et al., 2008).

Regardless of the power of using thousands of genome-wide SNPs, we did not detect fine scale genetic structure among wild populations. Instead, we found a seascape of genetic homogenization for wild populations in Peru and differentiation from the most distant population, in northern Chile, likely the result of long-distance dispersal limitations. The lack of population structure and outlier loci to distinguish wild populations limits the opportunity of using genomic tools to trace Peruvian scallop stocks back to their population of origin. However, the low heterozygosity observed in wild population nearby hatcheries suggest that genetic population descriptors such as kinship or inbreeding index could be used as an alternative to trace products back to wild stocks or to aquaculture hatcheries (i.e. bred in captivity).

## 5. CONCLUSION

We detected a genetic homogeneity among Peruvian populations and a strong differentiation with a population in the north of Chile. Our results suggest that the genetic similarity among Peruvian populations is a footprint of long-term man-made seed translocations, beyond what would be expected by larval dispersion and oceanographic factors. Our study indicates that the traceability of individuals from wild or aquaculture populations using molecular markers would be difficult, but we do not overrule the use of population descriptors, like kinship or inbreeding index, as an alternative.

## 6. ACKNOWLEDGEMENTS

The authors are grateful to Dr. Aldo Pacheco, who facilitated access to scallop samples from La Rinconada (Chile), and to Dr. Ronnie Gavilán and Dr. Omar Cáceres, from the Peruvian National Institute of Health for kindly facilitating the development of genomic libraries. This study was funded by grant No. 189-FINCyT-FIDECOM-PIPEA-2014 from The Fund for Innovation, Science and Technology (FINCyT/INNOVATE) of the Peruvian Ministry of Production.

## 8. AUTHOR CONTRIBUTIONS

Ximena Velez-Zuazo: Conceived and designed the project, wrote the research proposal, coordinated field data collection, analyzed the data, and wrote the study.

Sergio Barahona: Conducted laboratory analyses including DNA extractions and development of RAD libraries, analyzed the data, and wrote the study.

Omar G. Melo: Conducted laboratory analyses including DNA extractions and development of RAD libraries.

Eric Hanschke: Contributed to design of the project, co-wrote the research proposal, and helped collect field data,

Ian Hanschke: Contributed to design of the project, co-wrote the research proposal and helped collect field data.

Monica C. Santa-Maria: Conceived and designed the project, wrote the research proposal, coordinated laboratory analyses, and wrote the study.

