## Supplementary Data 1 for "Substantial gene flow caused by long-term translocation between wild populations of the Peruvian scallop (*Argopecten purpuratus*) is supported by RAD-Seq analyses"

SUPPLEMENTARY TABLE 1. RAD sequencing reads obtained with two runs of Illumina HiSeq 2500 for four libraries developed for five wild populations of *A. purpuratus*.

| Individual ID | Population | Barcode | Sequences | No. RadTag | Low Quality | Retained |
| --- | --- | --- | --- | --- | --- | --- |
| IND04 | B. Independencia | AACCC | 6346708 | 208586 | 36423 | 6101699 |
| IND05 | B. Independencia | CTGAA | 4461566 | 1669103 | 3272 | 2789191 |
| IND13 | B. Independencia | ACGTA | 2614037 | 71373 | 15257 | 2527407 |
| IND14 | B. Independencia | GCATT | 7087095 | 842608 | 58863 | 6185624 |
| IND16.1 | B. Independencia | GTACA | 1871775 | 108497 | 10465 | 1752813 |
| IND17 | B. Independencia | TCAGA | 4678319 | 600935 | 36686 | 4040698 |
| IND23 | B. Independencia | GTGTG | 4440545 | 560570 | 30307 | 3849668 |
| IND28 | B. Independencia | ACCAT | 3725540 | 368658 | 20825 | 3336057 |
| IND30 | B. Independencia | GTACA | 2608620 | 205839 | 22569 | 2380212 |
| LDT01 | Lobos de Tierra | AACCC | 6240286 | 263207 | 59381 | 5917698 |
| LDT05 | Lobos de Tierra | TAATG | 7354016 | 1054655 | 58162 | 6241199 |
| LDT09 | Lobos de Tierra | CGGCG | 655101 | 350491 | 2855 | 301755 |
| LDT10 | Lobos de Tierra | CAGTC | 2919011 | 216460 | 16604 | 2685947 |
| LDT11 | Lobos de Tierra | TGACC | 2396742 | 1166375 | 1440 | 1228927 |
| LDT15 | Lobos de Tierra | ACGTA | 3180193 | 144423 | 26854 | 3008916 |
| LDT16 | Lobos de Tierra | GCATT | 5477543 | 2885326 | 3002 | 2589215 |
| LDT18 | Lobos de Tierra | ACTGC | 498875 | 176639 | 2306 | 319930 |
| LDT21 | Lobos de Tierra | CGATA | 1283946 | 429588 | 5062 | 849296 |
| LDT24 | Lobos de Tierra | TGGTT | 3903879 | 664495 | 27715 | 3211669 |
| LDT25 | Lobos de Tierra | CTGAA | 5261094 | 266453 | 29448 | 4965193 |
| LDT26 | Lobos de Tierra | TGACC | 3089845 | 131111 | 17880 | 2940854 |
| RIN01 | Rinconada | GAAGC | 6064161 | 1724474 | 5017 | 4334670 |
| RIN02 | Rinconada | CAACT | 1572719 | 104953 | 8554 | 1459212 |
| RIN09 | Rinconada | CAACT | 3814317 | 206988 | 33316 | 3574013 |
| RIN10 | Rinconada | AATTT | 6571652 | 250865 | 39381 | 6281406 |
| RIN11 | Rinconada | CTAGG | 5797819 | 173462 | 32803 | 5591554 |
| RIN12 | Rinconada | AAGGG | 1802990 | 99950 | 9947 | 1693093 |
| RIN13 | Rinconada | TGGTT | 2368525 | 1240546 | 1287 | 1126692 |
| RIN14 | Rinconada | AAGGG | 2758098 | 240675 | 20576 | 2496847 |
| RIN15 | Rinconada | GCGCC | 7242993 | 368809 | 43500 | 6830684 |
| RIN17 | Rinconada | CTGAA | 4770866 | 616149 | 36221 | 4118496 |
| RIN19 | Rinconada | AATTT | 5101157 | 299147 | 43744 | 4758266 |
| RIN20 | Rinconada | CTAGG | 2622285 | 197151 | 19972 | 2405162 |
| SAM01 | Samanco | CCGGT | 2944527 | 181208 | 16095 | 2747224 |
| SAM02 | Samanco | GGAAG | 5255572 | 444592 | 40990 | 4769990 |
| SAM05 | Samanco | GAGAT | 8070413 | 1106779 | 62114 | 6901520 |
| SAM06 | Samanco | CCGGT | 4929936 | 428294 | 38444 | 4463198 |
| SAM08 | Samanco | GAAGC | 5749082 | 463575 | 54120 | 5231387 |
| SAM10 | Samanco | GGGGA | 3606386 | 249286 | 19555 | 3337545 |
| SAM12 | Samanco | TCAGA | 6485748 | 1221163 | 6053 | 5258532 |
| SAM18 | Samanco | TGACC | 5508834 | 354809 | 55263 | 5098762 |
| SAM24 | Samanco | GAAGC | 5603244 | 237172 | 30825 | 5335247 |
| SAM26 | Samanco | CCGGT | 10954065 | 1198676 | 11551 | 9743838 |
| SAM27 | Samanco | GAGAT | 8828125 | 576442 | 50216 | 8201467 |
| SAM28 | Samanco | TGGTT | 2673932 | 169929 | 14034 | 2489969 |
| SEC07 | Sechura | TAGCA | 4895777 | 165272 | 31019 | 4699486 |
| SEC09 | Sechura | GGGGA | 4492086 | 487317 | 33638 | 3971131 |
| SEC10 | Sechura | ACTGC | 3393593 | 241827 | 30136 | 3121630 |
| SEC12 | Sechura | CGGCG | 5806848 | 600822 | 60931 | 5145095 |
| SEC13 | Sechura | CGATA | 6053608 | 617977 | 49787 | 5385844 |
| SEC14 | Sechura | CCAAC | 3750821 | 333538 | 22863 | 3394420 |
| SEC16 | Sechura | GGAAG | 5378070 | 1820944 | 4254 | 3552872 |
| SEC17 | Sechura | GGAAG | 1028365 | 139254 | 5008 | 884103 |
| SEC20 | Sechura | CCAAC | 6257086 | 842344 | 64156 | 5350586 |
| SEC21 | Sechura | TAGCA | 5786839 | 196724 | 52813 | 5537302 |
| SEC26 | Sechura | CCAAC | 10670760 | 2059740 | 10257 | 8600763 |
| SEC27 | Sechura | TAATG | 910715 | 612698 | 3132 | 294885 |
