## Supplementary Data 2 for "Substantial gene flow caused by long-term translocation between wild populations of the Peruvian scallop (*Argopecten purpuratus*) is supported by RAD-Seq analyses"

SUPPLEMENTARY TABLE 2 Sequencing results for each population. N: Number of individuals. Sequences: Total number of sequences. Min: Minimum number of sequences per individual. Max: Maximum number of sequences per individual. SD: standard deviation.

| Population | N | Sequences | Min | Max | Average | SD |
| --- | --- | --- | --- | --- | --- | --- |
| Sechura | 12 | 58424568 | 910715 | 10670760 | 4868714 | 2575409.3 |
| Lobos de Tierra | 12 | 42260531 | 498875 | 7354016 | 3521711 | 2202653.9 |
| Samanco | 12 | 70609864 | 2673932 | 10954065 | 5884155 | 2434439.1 |
| B. Independencia | 9 | 37834205 | 1871775 | 7087095 | 4203801 | 1731766.6 |
| La Rinconada | 12 | 50487582 | 1572719 | 7242993 | 4207299 | 1971837.5 |
